# Translational regulation by bacterial small RNAs via an unusual Hfq-dependent mechanism

**DOI:** 10.1101/169359

**Authors:** Muhammad S. Azam, Carin K. Vanderpool

## Abstract

In bacteria, the canonical mechanism of translational repression by small RNAs (sRNAs) involves sRNA-mRNA base pairing that occludes the ribosome binding site (RBS), directly preventing translation. In this mechanism, the sRNA is the direct regulator, while the RNA chaperone Hfq plays a supporting role by stabilizing the sRNA. There are a few examples where the sRNA does not directly interfere with ribosome binding, yet translation of the target mRNA is still inhibited. Mechanistically, this non-canonical regulation by sRNAs is poorly understood. Our previous work demonstrated repression of the mannose transporter *manX* mRNA by the sRNA SgrS, but the regulatory mechanism was unknown. Here, we report that *manX* translation is controlled by a molecular role-reversal mechanism where Hfq, not the sRNA, is the direct repressor. Hfq binding adjacent to the *manX* RBS is required for sRNA-mediated translational repression. Translation of *manX* is also regulated by another sRNA, DicF, via the same non-canonical Hfq-dependent mechanism. Our results suggest that the sRNAs recruit Hfq to its binding site or stabilize the mRNA-Hfq complex. This work adds to the growing number of examples of diverse mechanisms of translational regulation by sRNAs in bacteria.

## INTRODUCTION

Almost 35 years ago, the first non-coding RNAs, RNA I and CopA, involved in plasmid copy number control, were discovered in bacteria (1,2). Since then, in all three domains of life, a surprisingly diverse yet poorly characterized set of regulatory RNAs has been discovered. Bacterial small RNAs (sRNAs), most in the size range of 50-250 nucleotides, act by imperfect, non-contiguous base pairing with mRNA targets to control translation or mRNA stability. There are also sRNAs that positively regulate targets, for example, by pairing with the 5′-UTR of an mRNA to prevent formation of a translation-inhibitory secondary structure (3,4) or pairing with ribonuclease recognition sequences in order to stabilize an mRNA target (5,6). For negative regulation, sRNAs often, but not always, operate as translational repressors by directly pairing with sequences overlapping the ribosome binding site (RBS), sequestering it from the incoming ribosome (7-9). To perform any of these regulatory tasks, sRNAs frequently depend on the chaperone protein Hfq. Hfq was initially discovered as the host factor for the replication of bacteriophage Qβ, but over the last few decades, its pleiotropic role in cellular physiology has reignited the interest of the research community (10). Hfq has emerged as a key factor in sRNA-mediated gene regulation, and in control of stability of mRNAs and sRNAs (11,12). Hfq is thought of as a matchmaking chaperone that binds to both sRNA and mRNA and promotes interaction between the sRNA and the target. Another key role of Hfq is protection of sRNAs from RNase E-mediated degradation (11,13,14). Hfq is a donut-shaped homohexameric protein belonging to the large family of Sm‐ and Sm-like proteins that are present in all three domains of life (11,15,16).

In this study, we uncovered an unconventional mechanism of regulation carried out by two Hfq-dependent sRNAs, SgrS and DicF, for translational repression of a target, *manX* mRNA. The physiological condition that triggers expression of DicF is unknown, but SgrS is expressed during glucose-phosphate stress (17). Sugars are critical nutrients that fuel central metabolic pathways to generate energy and precursor metabolites needed to synthesize nucleotides, amino acids, and fatty acids. Nonetheless, accumulation of excess phosphorylated sugar intermediates, and their non-metabolizable derivatives can be growth inhibitory (18,19). For instance, non-metabolizable sugar analogs, such as α-methylglucoside (αMG) or 2-deoxy-D-glucose (2DG), induce glucose-phosphate stress and production of SgrS (17,20). SgrS base pairs with mRNA targets to regulate their translation and stability (6,17,21,22). One of the key activities of SgrS during glucose-phosphate stress is repression of mRNAs encoding phosphotransferase system (PTS) sugar transporters, *ptsG* (17) and *manXYZ* (21,23). This repression inhibits new synthesis of PTS transporters and reduces uptake of sugars that are not efficiently metabolized during stress. We have shown that base pairing-dependent repression of transporter synthesis by SgrS is required for continued growth under stress conditions (24).

Many sRNAs that repress gene expression do so by inhibiting translation initiation by preventing ribosome binding to target mRNAs. Since bacterial translation initiation requires RNA-RNA base pairing between the 16S rRNA and the ribosome binding site (RBS), sRNAs typically base pair with a site close to the RBS and compete with the 30S ribosomal subunit (25). Initiating ribosomes occupy ~40 nucleotides (nts) on the mRNA, from −20 nt in the 5′ UTR to +19 nt into the coding region, where numbering is relative to the start codon (7,26). Work to define the region where a base pairing interaction could prevent formation of the translation initiation complex (TIC) indicated that base pairing within 15 nts downstream of the start codon can inhibit TIC formation (27). This led to the *“*five-codon window” hypothesis that proposed that if an sRNA base pairs with nucleotides comprising the first five codons of the mRNA, it can directly inhibit binding of the 30S ribosomal subunit and repress translation initiation. Interestingly, some studies have uncovered apparent exceptions to this hypothesis where sRNAs repress translation, either directly or indirectly, by base pairing outside of the five-codon window (22,28-30). For example, the Massé group found that binding of the sRNA Spot 42 at a site ~50 nt upstream of the *sdhC* start codon repressed *sdhC* translation. Their evidence suggested that the Spot 42 itself does not directly compete with the initiating ribosome, but instead may recruit the RNA chaperone Hfq to bind near the *sdhC* RBS to act as the primary repressor (28).

In this study, we investigated the mechanism by which SgrS regulates the first cistron of the *manXYZ* operon, *manX*. We observed previously that regulation of *manX* mRNA by SgrS involves base pairing 20 nt downstream of the start codon, which lies outside the 5-codon window (21). We also characterized regulation of *manX* translation by another sRNA regulator, DicF (31), a 53-nt long Hfq-dependent sRNA (32). The DicF binding site on *manX* mRNA is even further downstream from the start codon than the SgrS binding site. We hypothesized that each of these sRNAs regulates *manX* translation by a “non-canonical” mechanism, since their binding sites are positioned too far downstream for sRNA-mRNA base pairing to directly occlude ribosome binding. To test this hypothesis, we addressed several questions. Does sRNA-mRNA duplex formation directly inhibit translation by preventing formation of the translation initiation complex? If not, then is Hfq required for translational repression? Does Hfq bind to the *manX* mRNA near the ribosome binding site? Our results demonstrate that Hfq is absolutely required for translational repression mediated by SgrS and DicF in vivo. In vitro, Hfq, but not the sRNAs, can specifically inhibit formation of the TIC on *manX* mRNA. RNA footprints confirmed that SgrS and DicF have distinct binding sites in the *manX* coding region, and both sRNAs facilitate Hfq binding at a site close to the RBS. Taken together, our data demonstrate sRNAs mediate regulation of *manX* translation by a non-canonical mechanism involving recruitment or stabilization of Hfq binding at a site where it can directly interfere with translation.

## MATERIALS AND METHODS

### Strains and plasmids

The strains, plasmids, and oligonucleotides used in this study are listed in Supplementary Tables 1 and 2. Derivatives of *E. coli* K-12 MG1655 were used for all experiments. Alleles were moved between strains using P1 transduction (33) or λ-red recombination (34). Translational LacZ fusions, under the control of an arabinose-inducible P_BAD_ promoter, were constructed by PCR amplifying DNA fragments using primers with 5′-homologies with the promoter and to the 9^th^ codon of *lacZ* (Table S2). These fragments were integrated into the chromosome by λ-red recombination using counterselection against *sacB* as described previously (35).

SA1328, a strain with tet-Cp19-115nt-*manX'-'lacZ, ΔsgrS, lacI^q^,* kan^R^, Δ*hfq* genotype was constructed in two steps. First, Δ*hfq::FRT-kan-FRT* was transduced into JH111 and pCP20 was used to flip out the kan^R^ cassette. The tet-Cp19-115nt-*manX'-'lacZ* cassette was then transduced into the latter strain.

Strains containing truncated *manX* translational fusions under the control of a P_BAD_ promoter were constructed in strain PM1205 (35). The P_BAD_-22nt-*manX'-'lacZ* and P_BAD_-25nt-*manX'-'lacZ* fusions were generated by PCR amplifying DNA fragments with primer pairs O-SA178/O-SA176 and O-SA177/O-SA176 primer pairs respectively, containing 5′ homologies to pBAD and *lacZ* (Table S2). The PCR products were recombined into PM1205 using λ-red homologous recombination as described previously. The same fusions with mutations in the Hfq binding site, mut-1, (in strains SA1522 and SA1620), were created using the method above, but using oligonucleotides O-SA177/O-SA176 and O-SA177/O-SA433 to obtain the PCR products. Oligonucleotide pairs O-SA645/O-SA176, O-SA646/O-SA176, O-SA647/O-SA176, and O-SA648/O-SA176 to obtain PCR products with mut-2, mut-3, mut-4, and mut-5 binding site mutations, respectively.

### Media and regents

Unless otherwise stated, bacteria were cultured in LB broth or on LB agar plates at 37°C. TB medium was used for β-Galactosidase assays. A final concentration of 0.1 mM IPTG (isopropyl β-D-1-thiogalactopyranoside) was used to induce Lac promoters. L-arabinose was used at concentrations of 0.001%, for solid media, and 0.002%, for liquid media, to induce P_BAD_ promoters. Antibiotics were used at following concentrations: 100 µg/ml ampicillin, 25 µg/ml chloramphenicol, 50 µg/ml spectinomycin, and 25 µg/ml kanamycin.

### β-Galactosidase assays

Strains with *lacZ* fusions were grown overnight in TB medium and subcultured 1:100 to a fresh medium containing Amp and 0.002% L-arabinose (for P_BAD_ promoters). Cultures were grown at 37°C with shaking to OD_600_~0.2. At this point, sRNAs were induced with 0.1 mM IPTG (final concentration) and cells were grown for another hour to OD_600_~0.5. β-Galactosidase assays were performed on these cells according to the previously published protocol (36). The error bars represent standard deviation derived from three biological replicates.

### In vitro transcription

For in vitro transcription, template DNA was generated by PCR using gene specific oligonucleotides with a T7 promoter sequence at the 5′ end of the forward primer. The following oligonucleotides were used to generate templates for RNA footprinting and gel shift assays: O-JH219/O-JH119 and O-JH218/O-JH169 to generate *manX* and SgrS template DNA. DNA template for DicF transcription was generated by hybridizing two oligos, DicFW and DicFC, in TE buffer. Transcription of these DNA templates was performed using the MEGAscript T7 kit (Ambion) following manufacturer's instructions.

### Purification of His-tagged Hfq

Hfq-His protein was purified following a previously published protocol (37). BL21(DE3) cells harboring pET21b-Hfq-His_6_ was cultured in 400 mL LB medium at 37°C. At OD_600_~0.3, IPTG was added to a final concentration of 1 mM, and the incubation was continued for 2 hrs. The cells were washed with STE buffer (100 mM NaCl; 10 mM Tris · HCl, pH 8.0; 1 mM EDTA) and resuspended in 10 mL Equilibration buffer (50 mM Na_2_HPO_4_–NaH_2_PO_4_, 300 mM NaCl, and 10 mM imidazole). The suspension was treated with 25 mg lysozyme, incubated on ice for 10 minutes and sonicated. The supernatant was collected after centrifugation at 16,000 × g for 10 min at 4°C followed by incubation at 80°C for 10 minutes. The sample was centrifuged again, at 16,000 × g for 10 min at 4°C. The supernatant was fractionated using a Ni^2+^NTA agarose column following manufacturer's instructions (Roche) and checked by SDS-PAGE electrophoresis. The fractions containing Hfq were pooled, dialyzed, and stored in a storage buffer (20 mM Tris · HCl pH 8.0, 0.1 M KCl; 5 mM MgCl_2_, 50% glycerol, 0.1% Tween 20, and 1 mM DTT) at −20°C.

### Toeprinting assays

Toeprinting assays were performed using unlabeled *manX_ATG_* and P^32^-end-labeled primer in the presence and absence of Hfq and SgrS following the previously published protocol (38). For each reaction, 2 pmol of *manX* RNA and 1.6 pmol of end-labeled primer (O-JH119) were heated for one min at 95°C in toeprint buffer (20 mM Tris-HCl pH 7.5, 50 mM KCl, 0.1 mM EDTA, and 1mM DTT). The mixture was chilled in ice for five minutes, followed by addition of MgCl_2_ and dNTPs (10 mM and 1 mM respectively, final concentrations). Purified Hfq and in vitro synthesized SgrS RNA were also added to the appropriate reactions and incubated at 37°C for 10 minutes. Next, ribosomes (1.3 pmol, NEB) were added to this reaction mixture, and the incubation was continued at 37°C for five minutes. Thirteen picomoles of fMet-tRNA (Sigma) was added to this reaction and cDNAs were synthesized using SuperScript III reverse transcriptase (Invitrogen). The reaction was stopped by adding 10 µl of loading buffer II (Ambion). The reaction products were analyzed on an 8% polyacrylamide-urea gel. Sequencing ladders were generated using Sequenase 2.0 DNA sequencing kit (Affymetrix).

### Footprinting assays

In vitro RNA footprinting reactions were performed as described previously (39) with some modifications. 0.1 pmol of 5′-end labeled *manX* mRNA was incubated at 37°C for 30 minutes in structure buffer (Ambion) containing 1 ng of yeast RNA (Ambion), in the presence or absence of 78 pmoles of unlabeled SgrS, 240 pmoles of unlabeled DicF, and 3.7 pmoles of Hfq. At this point, lead acetate (Sigma) was added to a final concentration of 2.5 µM for the cleavage reaction and incubated at 37°C for two minutes. Reactions were stopped by adding 12 µL of loading buffer (Ambion). A modified protocol was followed to investigate Hfq binding to *manX* mRNA in the absence and presence of SgrS with limiting Hfq concentrations. To perform this footprint experiment, we used 2 ng of yeast RNA (Ambion) in the structure buffer, and 0.31 pmoles of Hfq was added to the indicated reactions. The alkaline ladder was generated by incubating 5′-end labeled *manX* mRNA at 90°C for 5 minutes in alkaline buffer (Ambion). RNase T1 was used for 5 minutes at 37°C to generate the G ladder. The samples were resolved on an 8% polyacrylamide/urea gel.

### Electrophoretic mobility shift assay

RNA-RNA and RNA-protein gel electrophoretic mobility shift assays were performed by incubating 0.01 pmol of P^32^-labeled denatured *manX* RNA with indicated amounts of SgrS or DiF (both denatured at 95°C for 1 min) or Hfq in binding buffer (20mM Tris-HCl pH 8.0, 1mM DTT, 1mM MgCl_2_, 20mM KCl, 10mM Na_2_HPO_4_–NaH_2_PO_4_ pH 8.0). The mixture was incubated at 37°C for 30 minutes, and non-denaturing loading buffer (50% glycerol and 0.1% bromophenol blue) was added. The samples were resolved on a 4.6% native polyacrylamide gel for 1.5 hours at 10 mA. The fraction of *manX* RNA bound was determined using Fluorescent Image Analyzer FLA-3000 (FUJIFILM) to quantitate the intensities of the bands. The data were fit into Sigmaplot software to obtain the *K*_D_ value.

## RESULTS

### *Hfq is essential for translational repression of* manX

Previously, we identified *manXYZ* mRNA, which encodes a mannose and (secondary) glucose transporter (EII^Man^), as a target of SgrS (21,23). The polycistronic *manXYZ* mRNA is negatively regulated post-transcriptionally via two independent SgrS-*manXYZ* mRNA base pairing interactions (23). The physiological outcome of this regulation is repression of EII^Man^ transporter synthesis, which helps rescue cell growth during glucose-phosphate stress (21). One of the SgrS binding sites was mapped to the coding region of *manX*, and this binding site was shown to be necessary for translational repression of *manX* (21,23). Although we have identified the base pairing sites for SgrS on the *manXYZ* transcript and established that translational regulation of this target is important for cell growth during glucose-phosphate stress (24), the exact mechanism of SgrS-mediated *manX* translational repression is unknown. The SgrS binding sites on the *manXYZ* mRNA are too far from the RBS of the *manX* and *manY* cistrons (23) to directly occlude ribosome binding. We hypothesized that *manX* translation might be repressed by a non-canonical mechanism where Hfq serves as the direct repressor of translation while SgrS plays an accessory role, perhaps recruiting Hfq to bind stably near the ribosome binding site (RBS). To test this hypothesis, we first investigated whether translational repression of two different SgrS targets, *ptsG* and *manX,* was dependent on Hfq in vivo. Aiba and coworkers showed that SgrS can inhibit translation of *ptsG* in the absence of Hfq (37), consistent with the canonical model for repression where the sRNA pairs near the translation initiation region (TIR) (Figs. S1, S2) and directly occludes ribosomes. Here, we utilized *manX'-'lacZ* and *ptsG'-'lacZ* translational fusions that we demonstrated are good reporters for SgrS-dependent regulation of these targets (21). We monitored fusion activities after SgrS production was induced in wild-type and Δ*hfq* backgrounds. Since stability of *E. coli* SgrS (SgrS*_Eco_*) is greatly reduced in the absence of Hfq (40), we utilized the *Salmonella* SgrS allele (SgrS*_Sal_*) which is more stable than SgrS*_Eco_* in the Δ*hfq* background. SgrS*_Sal_* has a very similar seed (Fig. S3) that displays comparable regulation of SgrS*_Eco_* targets (41). In agreement with our previous study (41), we found that SgrS*_Sal_* can complement a Δ*sgrS* mutant for regulation of *manX* (Fig. 1A). Consistent with the direct ribosome occlusion mechanism for translational regulation of *ptsG,* SgrS*_Sal_* efficiently repressed the *ptsG'-'lacZ* fusion in both the wild-type and Δ*hfq* backgrounds (Fig. 1B). In contrast, while SgrS efficiently repressed *manX* in a wild-type background, it failed to repress the fusion in an *hfq* mutant background, even at high levels of inducer (Fig. 1C). These data indicate that SgrS cannot regulate *manX* in vivo in the absence of Hfq.

**Figure 1.**
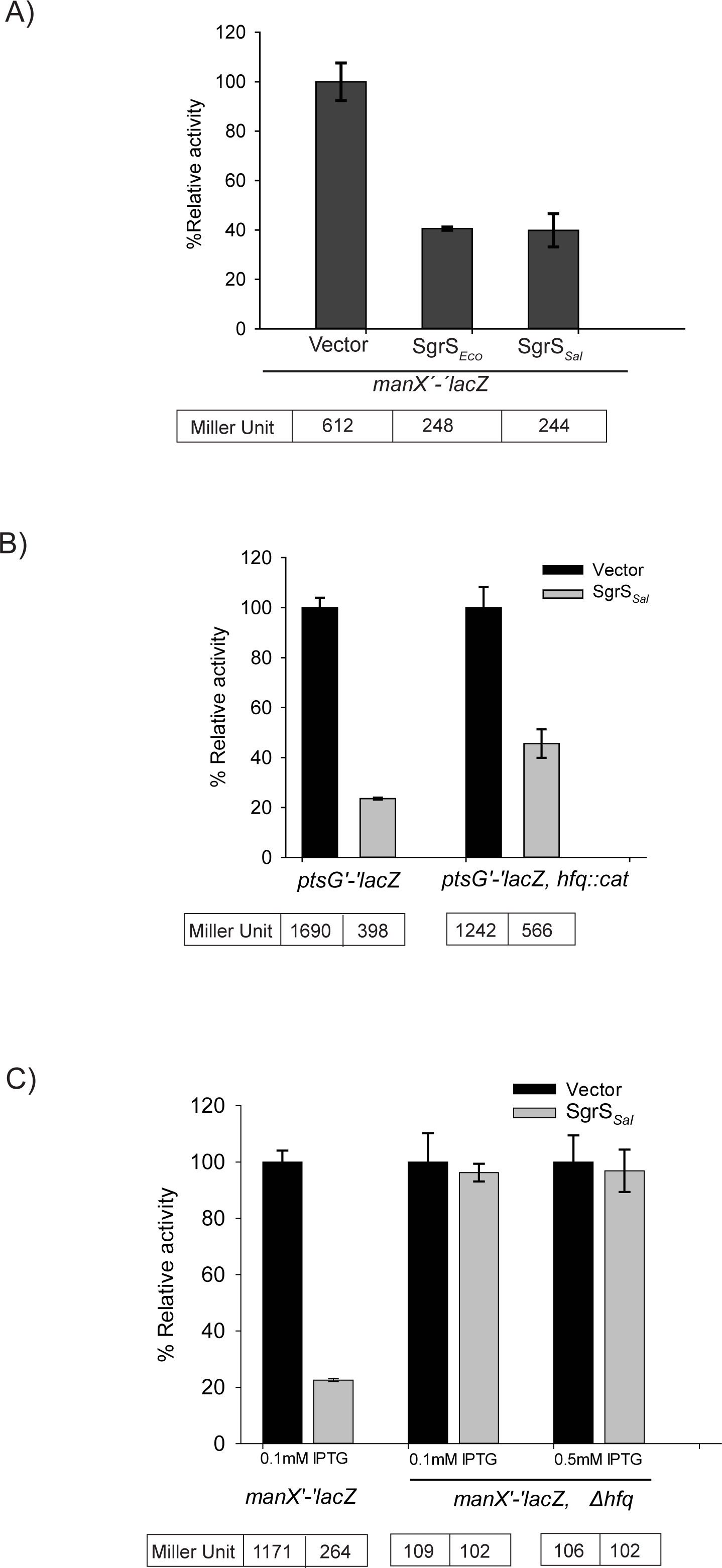
SgrS-mediated translational repression of *manX* is Hfq-dependent. **A)** Strain JH175 has a *manX'-'lacZ* translational fusion under the control of Cp19, a constitutive promoter. This strain was transformed with a vector control or plasmids with SgrS homologs from *Salmonella* (SgrS*_Sal_*) or *E. coli* (SgrS*_Eco_*), expressed after induction with 0.1 mM IPTG, and assayed for β-galactosidase activity after 60 min. Units of activity in the experimental samples were normalized to the levels in the vector control strains to yield percent relative activity. **B)** The same vector control and SgrS*_Sal_* plasmids described in A were transformed to strain JH184 (*hfq*+) or DB151 (Δ*hfq*), containing a Cp19-*ptsG'-'lacZ* fusion. Induction of SgrS and β-Galactosidase assays were conducted and analyzed as in A. **C)** Strains JH175 (*hfq*+) and SA1328 (Δ*hfq*) with the Cp19-*manX'-'lacZ* fusion were transformed with vector control or SgrS*_Sal_* plasmids. SgrS was induced using the indicated concentrations of IPTG and β-Galactosidase assays were conducted and analyzed as in A.

We hypothesized that the role of SgrS in *manX* regulation is to recruit or enhance binding by Hfq to the TIR. To identify a putative Hfq binding site in the 5′ UTR required for translational regulation, we constructed a series of *manX* translational fusions with truncations in the 5′ UTR (Fig. 2A). Activity of these fusions was measured in the presence and absence of ectopically expressed *sgrS*. The constructs contained 65, 30, 25, and 22 nt of the *manX* 5′ UTR, and in the *hfq^+^* background, all four fusions showed a similar pattern of regulation compared to the construct containing the full-length 115-nt *manX* 5' UTR (Fig. 1A, 2A). Further truncation of the *manX* 5' UTR was not possible without interfering with the RBS. The fact that all truncated fusions were regulated similarly to full-length suggested that the putative Hfq binding site resided downstream of the 5' boundary defined by the 22-nt fusion (Fig. 2A). An A/U-rich motif just upstream of the *manX* RBS is similar to the motif that was shown in other studies to be preferentially bound by Hfq (42,43). To test the role of this motif in the regulation of *manX* translation, we constructed a mutant *manX* fusion where the A/U-rich motif was converted to a G/C-rich motif (Fig. 2B). The basal level of *manX'-'lacZ* activity was dramatically reduced by the mutation (*mut-1*), perhaps because the motif is directly adjacent to the *manX* RBS and the mutation diminishes translational efficiency. Nevertheless, in contrast with the wild-type *manX* fusion, which was efficiently repressed when SgrS was ectopically expressed, SgrS failed to alter translation of the *mut-1* fusion (Fig. 2B). Importantly, *mut-1* is located ~35 nt upstream of the known SgrS binding site on *manX.* Loss of SgrS-dependent regulation caused by this mutation supports the notion that SgrS is not the direct regulator of *manX* translation.

**Figure 2.**
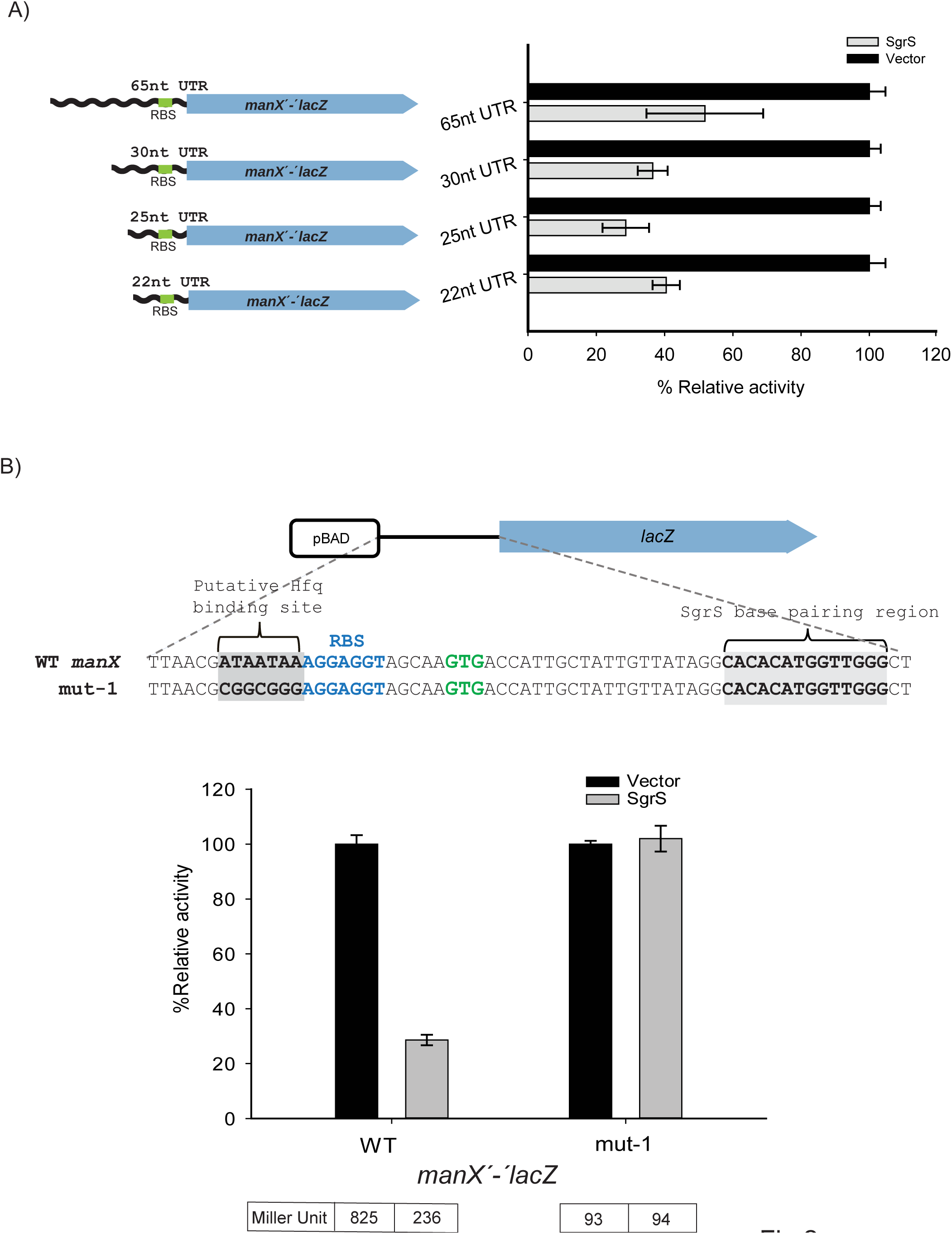
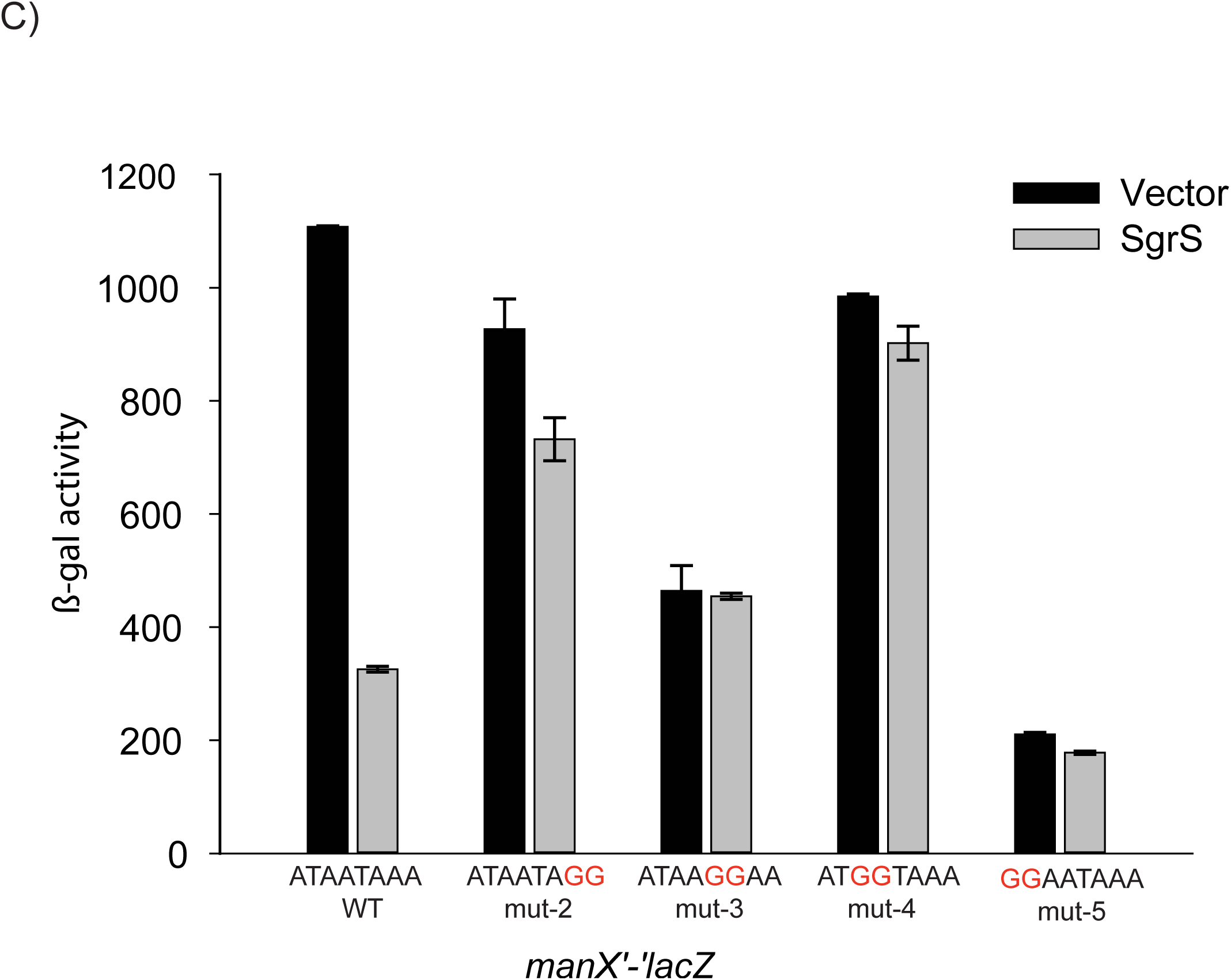
Genetic analysis of a putative Hfq binding site in the *manX* 5' UTR. **A)** The full-length *manX* UTR in *E. coli* is 115-nt. Four translation fusions with truncations of the UTR (as indicated) were constructed by moving the heterologous promoter closer to the TIR. Vector control and SgrS plasmids were transformed into the resulting strains (JH178, JH181, SA1404 and SA1403) and β-Galactosidase assays were conducted and analyzed as described for Fig. 1A. **B)** An A/U rich motif upstream of the *manX* RBS, was mutated in the context of the 25-nt *manX* translational fusion, resulting in the mut-1 fusion. The positions of the putative Hfq binding site and confirmed SgrS binding site are indicated with gray boxes. The RBS is in blue letters and the *manX* start codon is in green. The activity of SgrS on wild-type (strain SA1404) and mut-1 (strain SA1522) fusions was assessed after induction as described for Fig. 1A. **C)** Additional mutations as indicated in red were constructed in the putative Hfq binding site in the *manX* translational fusion, resulting in mutant fusions mut-2 through mut-5 (strains SA1713, SA1711, SA1712, SA1710, respectively). The plasmids, induction and β-Galactosidase assays were conducted and analyzed as described for Fig. 1A, except that activities are reported in Miller Units.

To further test the importance of nucleotides in the A/U rich motif adjacent to the *manX* RBS for SgrS‐ and Hfq-dependent regulation, we constructed four additional strains, each harboring 2-nt mutations (*mut-2-mut-5,* Fig. 2C). Compared to the wild-type *manX'-'lacZ* fusion, these mutants had variable basal levels of activity (Fig. 2C, vector), implying that the nt in this region play a role in overall translational efficiency. Nonetheless, all of the mutations caused loss of SgrS-mediated regulation (Fig. 2C, SgrS, and 3A). The *mut-4* fusion was particularly informative, as it retained nearly wild-type basal levels of activity, and showed a nearly complete loss of regulation by SgrS. These results suggest that this 2-nt mutation, changing the wild-type mRNA sequence from “AUAAUAAA” to the *mut-4* sequence “AUGGUAAA” was sufficient to disrupt SgrS-mediated regulation of *manX*, translation, possibly by preventing Hfq binding at this site.

The initial experiment suggesting that SgrS-dependent regulation of *manX,* but not *ptsG,* requires Hfq also revealed a striking reduction in basal levels of *manX'-'lacZ* activity in the *hfq* mutant (Fig. 1C, 109 Miller Units) compared to the wild-type background (Fig. 1C, 1170 Miller Units). To investigate the link between the putative Hfq binding site upstream of the *manX* RBS and *manX* translation and mRNA stability, we compared activities of wild-type and mutant fusions (*mut-1-mut-5*) in *hfq^+^* and Δ*hfq* backgrounds. We again saw that the basal levels of activity varied widely among mutant fusions, but for wild-type and all mutant fusions, activity was strongly reduced in the Δ*hfq* compared to *hfq^+^* background (Fig. 3A). These results suggested that differences in basal levels of activity of the mutant fusions were not due specifically to Hfq binding at the site adjacent to the *manX* RBS, since mutation of *hfq* further reduced activity of the mutant fusions (Fig. 3A). We reasoned that if Hfq binding, perhaps at another site, protects *manX* mRNA from RNase E-mediated degradation, then loss of Hfq could result in increased *manX* mRNA turnover and lower levels of *manX'-'lacZ* activity. To test this, we monitored activity of the *manX'-‘lacZ* fusions in an *rne131* background, which produces a truncated RNase E that cannot form the degradosome complex (44). Consistent with our prediction, in the *rne131* strain, *manX'-'lacZ* activity was approximately equivalent in *hfq^+^* and Δ*hfq* backgrounds for wild-type and all mutant fusions (Fig. 3B). To confirm that the *rne131* mutation impacted *manX* mRNA stability independent of any effect on SgrS-dependent regulation via Hfq binding to the site adjacent to the RBS, we conducted a control experiment (Fig. S4). In this experiment, we saw that a wild-type *manX'-'lacZ* fusion was regulated by SgrS as expected in the *rne131* mutant background whereas the mut-2 fusion (with the mutation in the Hfq binding site as in Fig. 2C) was not repressed by SgrS.

**Figure 3.**
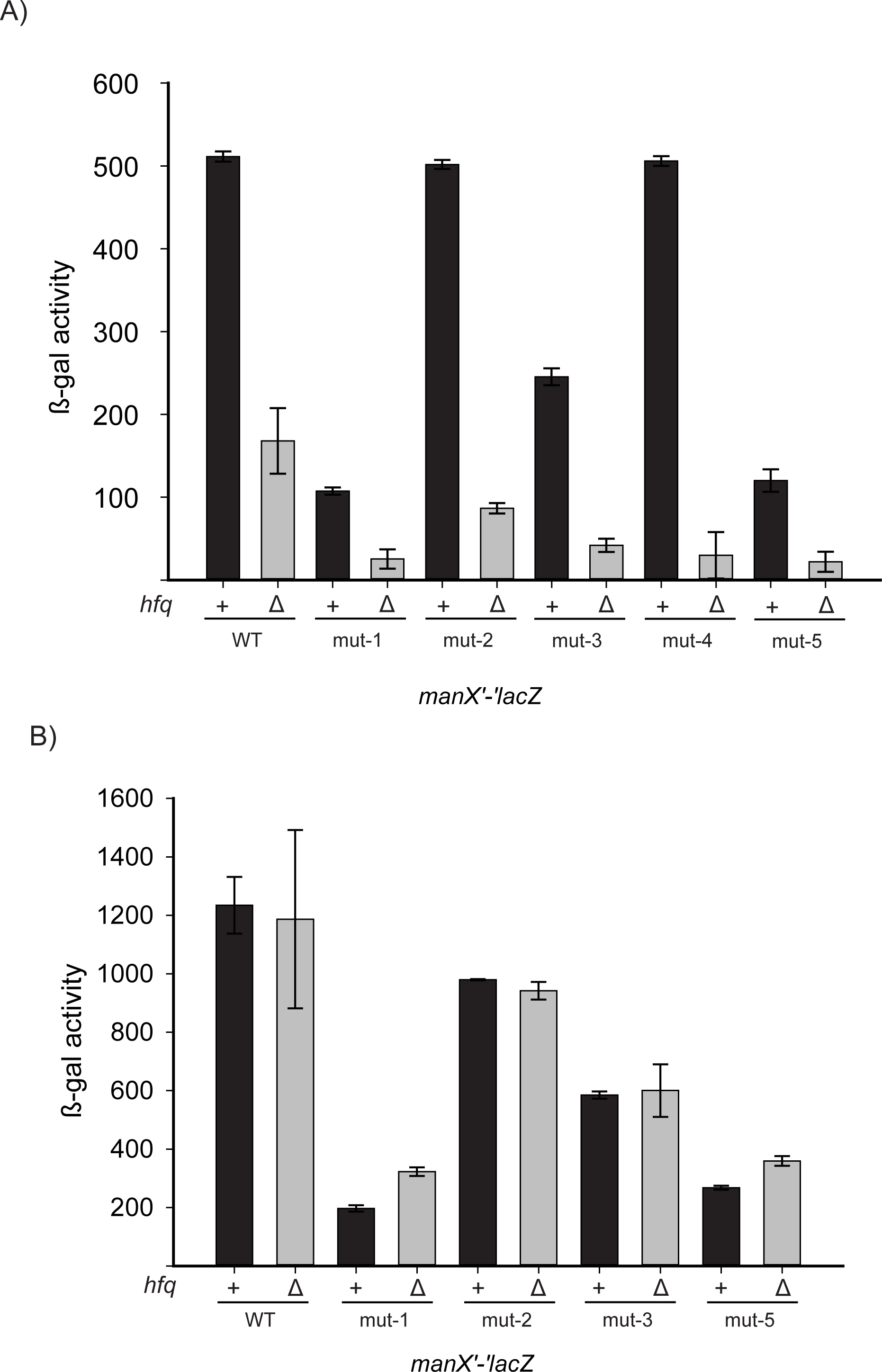
Mutations Hfq impact translation and stability of *manX* mRNA. **A)** Wild-type and mut-1 through mut-5 *manX* translational fusions (same as Fig. 2C) in *hfq^+^* (+) and Δ*hfq* (Δ) backgrounds (for strains, see Table S1) were assayed for β-Galactosidase activity. Cultures were grown to mid-log phase and then assayed. Specific activity is reported in Miller Units. **B)** The *rne131* allele was moved into the reporter strains described in A. β-Galactosidase activity was assayed as described in A.

Considering the results presented so far, we can conclude several points regarding *manX* regulation. First, SgrS alone is not sufficient for translational repression of *manX,* but requires Hfq for this activity (Fig. 1C). A putative Hfq binding site adjacent to the *manX* RBS, far upstream of the SgrS binding site, is required for SgrS-dependent repression of *manX* (Fig. 2B and 2C). Changes in basal (SgrS-independent) activity of *manX* fusions revealed that changes in sequences adjacent to the *manX* RBS alter translational efficiency (Fig. 2C). Furthermore, Hfq plays a role in protecting the *manX* mRNA from RNase E-mediated degradation, because reductions in *manX'-'lacZ* activity caused by deletion of *hfq* (Fig. 3A) were lost in an *rne131* background (Fig. 3B). Since this this occurred in wild-type and *mut-1-5* fusion strains, we suggest that Hfq stabilizing *manX'-'lacZ* mRNA involves an Hfq binding site different from the one involved in SgrS-dependent *manX* regulation. While this complex regulation at the levels of translation and mRNA stability is intriguing and warrants further investigation, we focus in this study on understanding how sRNA-mediated repression of *manX* translation via Hfq occurs.

### *Hfq inhibits formation of the translation initiation complex on* manX *mRNA*

If Hfq is the direct repressor of *manX* translation, it should compete with the ribosome for binding to the mRNA in vitro. We used toeprinting assays (38) to test whether Hfq or SgrS could directly inhibit translation initiation. In a toeprinting assay, stable binding of the 30S ribosomal subunit and tRNA^fMet^ to the RBS blocks a primer extension reaction and produces a product with a characteristic size. Since native *manX* has a weak GTG start codon that does not stably associate with commercially available preparations of 30S ribosomes, we changed the start codon to the canonical ATG to ensure strong initiation complex formation *in vitro*. We showed previously that this construct, *manX*_ATG_, was efficiently repressed by SgrS (21). The toeprint assay was performed by mixing *manX*_ATG_ mRNA, P^32^ end-labeled primer, ribosomes, and tRNA^fMet^ in the presence and absence of Hfq. Reverse transcriptase was then added to begin the primer extension reaction. In the positive control reaction, we saw the characteristic toeprint signal caused by termination of reverse transcription at position +15/+16 relative to the start codon (Fig. 4A). With the addition of increasing concentrations of Hfq, the formation of the TIC was completely inhibited (Fig. 4A and 4B). However, when increasing concentrations of SgrS were added in the absence of Hfq, TIC formation was unperturbed (Fig. 4B). When the same concentrations of SgrS were added to toeprint reactions with *ptsG* mRNA, we observed inhibition of the toeprint signal (Fig. S5). These results are consistent with the in vivo studies, and add further evidence supporting the hypothesis that SgrS-mediated regulation of *manX* occurs by a fundamentally different mechanism than *ptsG.* The data suggest that Hfq itself directly inhibits *manX* translation at the initiation stage.

**Figure 4.**
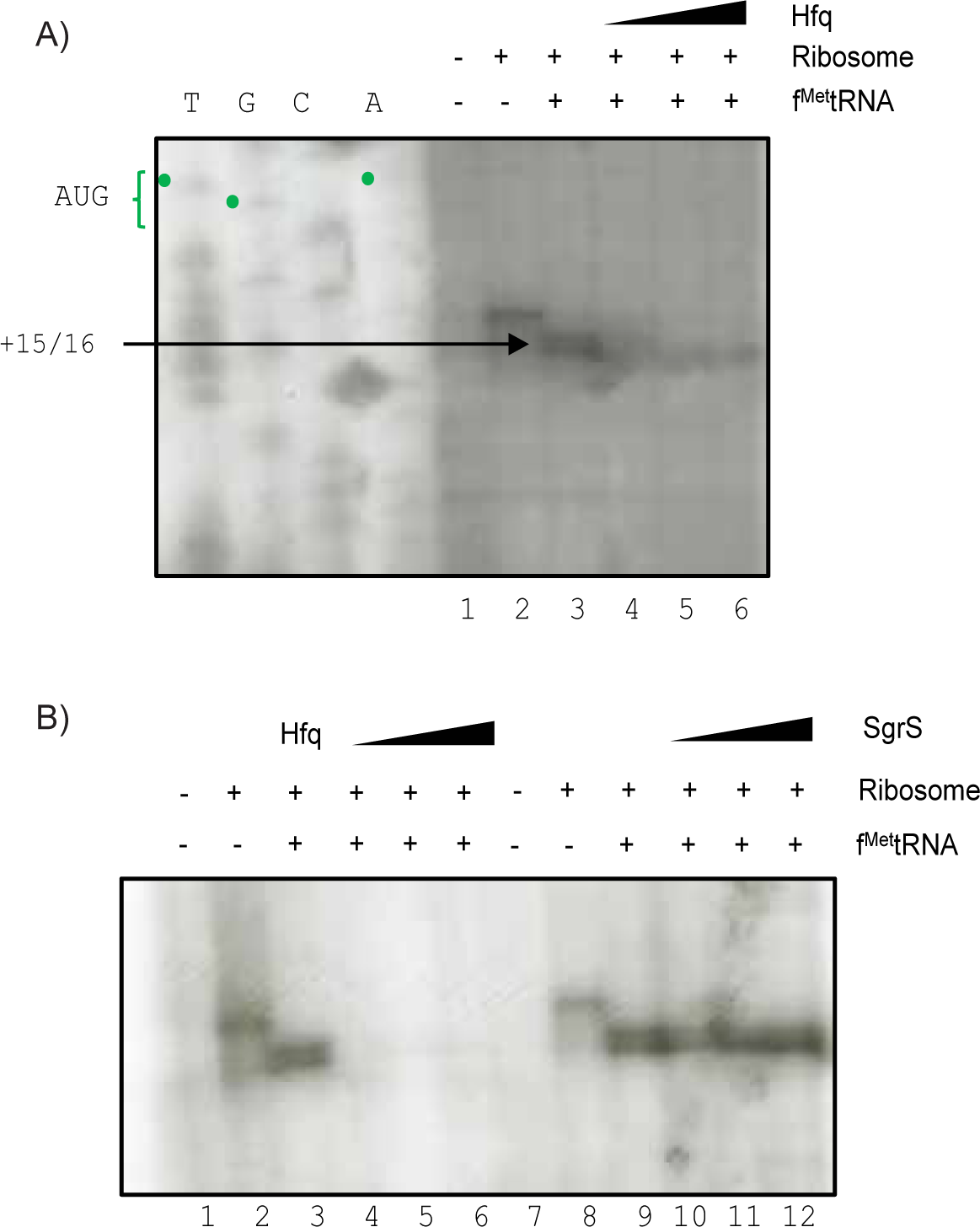
Toeprint assays reveal that Hfq, but not SgrS, can prevent ribosome binding to *manX* mRNA. **A)** Toeprint assays were conducted as described in Materials and Methods. Ribosomes, tRNA^fMet^ and Hfq were added to *manX* mRNA as indicated above the gel image. In lanes 4-6, Hfq concentrations were 0.15 µM, 0.5 µM, and 1 µM, respectively. The sequencing ladder is indicated by “T – G – C – A”, and was generated with the same oligo (OJH119) used for reverse transcription. The toeprint signal is indicated at +15/16 relative to the start codon. **B)** Lanes 1-6 on the left represent the same reactions as described in part A. Lanes 1-6 on the right represent similar reactions, except SgrS was added at concentrations of 100 nM, 250 nM, and 500 nM (lanes 10-12).

### manX *is regulated by DicF sRNA*

In a previous study, *manX* was identified as a putative target of another sRNA, DicF (31). To further investigate the regulation of *manX* by DicF and determine the regulatory mechanism, we monitored the activity of a *manX'-'lacZ* translational fusion (under the control of a constitutive promoter to rule out indirect effects on *manX* transcription) in control cells and cells where DicF was ectopically expressed. Cells expressing *dicF* showed ~40% reduced β-galactosidase activity compared to control cells (Fig. 5A). Compared to SgrS, which reduces *manX* translation by about 70% (Fig. 2B), DicF is a rather weak regulator.

**Figure 5.**
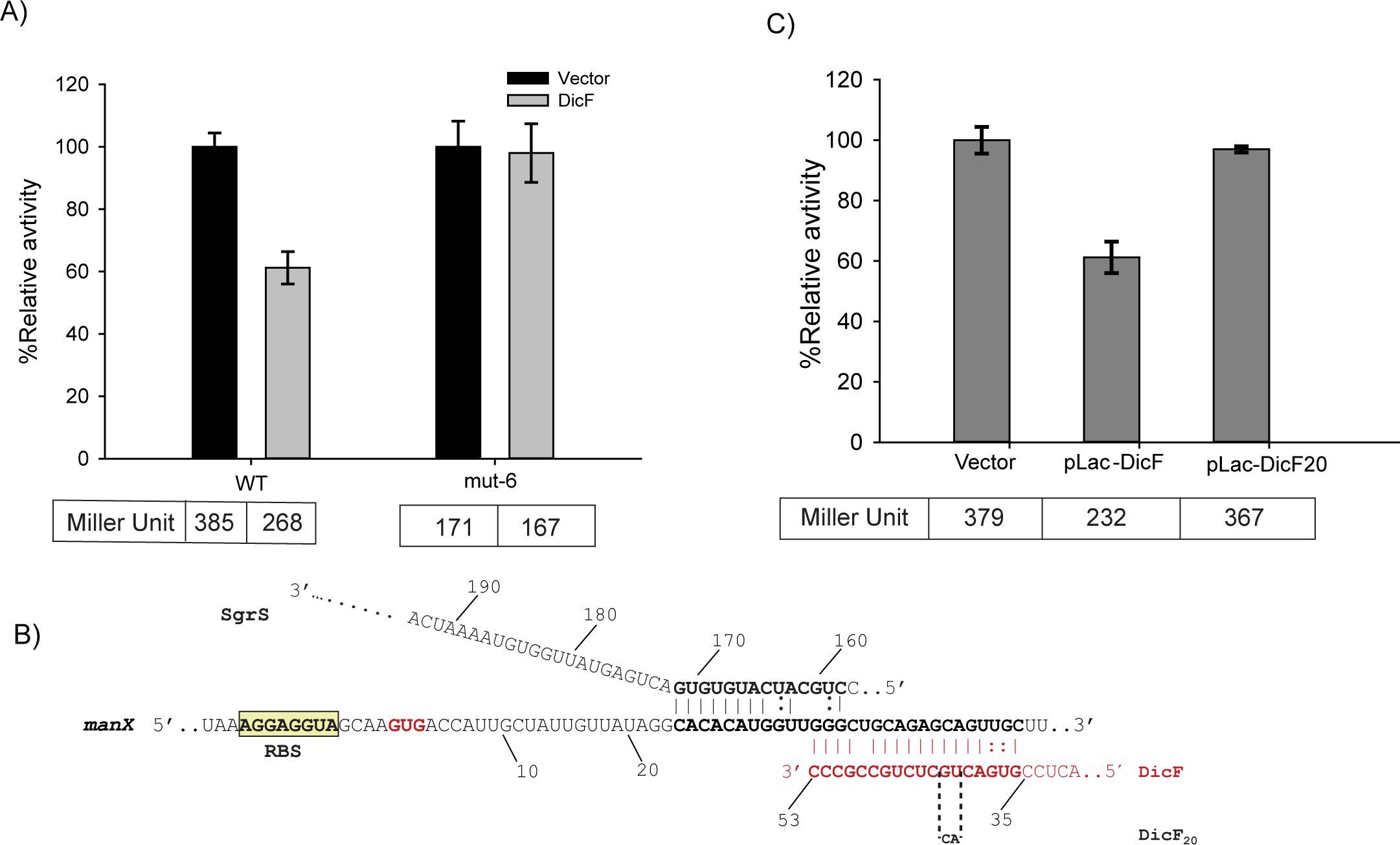
DicF, a prophage-encoded sRNA, also regulates *manX* translation. **A)** Strains with *manX'-'lacZ* fusions with a wild-type (JH175) or mutant (putative) Hfq binding site mut-6 (SA1620) (as shown in Fig. 2B) were transformed with vector control or DicF-expressing plasmids. Expression of DicF was induced with 0.1 mM IPTG, and β-Galactosidase assays were conducted and analyzed as described for Fig. 1A. **B)** Base pairing interactions for *manX* mRNA (middle sequence) and SgrS (top sequence) or DicF (bottom sequence). The position of the DicF20 mutation is indicated below the DicF sequence. **C)** Strain JH175, containing the wild-type *manX'-‘lacZ* fusion, was transformed with vector control or plasmids expressing wild-type DicF or mutant DicF20. Expression of DicF was induced with 0.1 mM IPTG, and β-Galactosidase assays were conducted and analyzed as described for Fig. 1A.

Previous studies have demonstrated that sequences at either the 5′ or the 3′ end of DicF can base pair with mRNA targets (31,45). We identified a potential base pairing interaction between the 3′ end of DicF and the coding region of *manX* just downstream of the known SgrS binding site (Fig. 5B). To test this base pairing prediction, we made a mutation in nucleotides of DicF that should disrupt the base pairing interaction (Fig. 5B). This mutant allele, *dicF20*, lost the ability to regulate the *manX'-'lacZ* translational fusion (Fig. 5C), consistent with the base pairing prediction. If DicF also base pairs within the *manX* coding sequence, well outside the window that would allow direct interference with ribosome binding, then like SgrS, DicF may also repress *manX* translation by influencing Hfq binding in the *manX* TIR. To test whether DicF-mediated regulation requires the putative Hfq binding site near the RBS, we constructed a mutant version of the *manX'-'lacZ* fusion (containing the putative DicF base pairing site) where the A/U-rich region next to the RBS is changed to G/C-rich (*mut-6*). In contrast with the wild-type *manX* fusion, which was repressed upon *dicF* expression, activity of the *mut-6* fusion with the mutation in the putative Hfq binding site was not altered by DicF (Fig. 5A). This observation is consistent with the model that DicF-mediated regulation of *manX* also requires Hfq binding proximal to the RBS where it acts as the direct translational repressor.

### *Hfq binds next to the* manX *ribosome binding site*

We predicted that DicF base pairs at a site just downstream of the SgrS binding site, from residues G145 to C162 (Fig. 5B). Our genetic analyses suggest that Hfq binds in the 5′ UTR just upstream of the RBS to act as the direct repressor of *manX* translation for both SgrS‐ and DicF-mediated regulation (Figs. 2B and 5A). To further test these predictions, we performed in vitro footprinting experiments with labeled *manX* mRNA to identify the Hfq binding site(s) occupied in the presence of each individual sRNA. As we showed previously, SgrS protects its binding site from C139 to G152 on *manX* mRNA even in the absence of Hfq, and the addition of Hfq does not change the footprint (Fig. 6A) (21). Notably, SgrS alone does not affect the structure around the RBS or start codon. Consistent with our prediction (Fig. 6B), DicF protects *manX* mRNA from G150 to C167 in the absence and presence of Hfq. Again, DicF only impacted the reactivity of nucleotides comprising its binding site in the *manX* coding region, and the structure upstream in the TIR was unaffected. In the presence of either sRNA, Hfq clearly protected *manX* mRNA nucleotides A97-A103 (Fig. 6A). Note that this region is the same A/U-rich region that we predicted as the Hfq binding site (Fig. 2B) and that when mutated, prevented SgrS‐ and DicF-dependent regulation (Figs. 2B and 5A, respectively). Additionally, we observe some weak protection of residues U91-C95. These findings demonstrate that Hfq binds at the same location on *manX* mRNA, adjacent to the RBS, regardless of which sRNA is present in the sRNA-mRNA-Hfq ternary complex.

**Figure 6.**
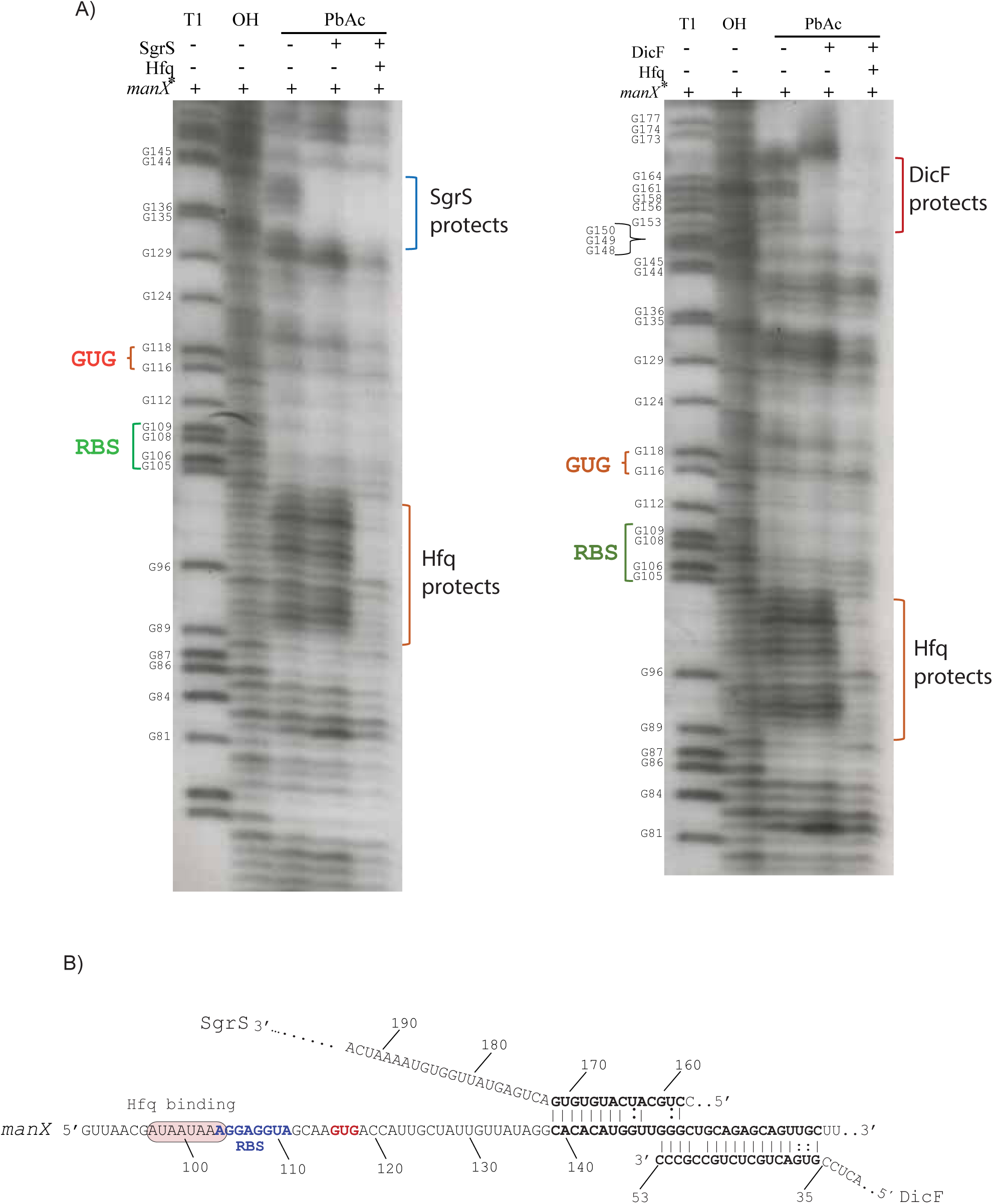
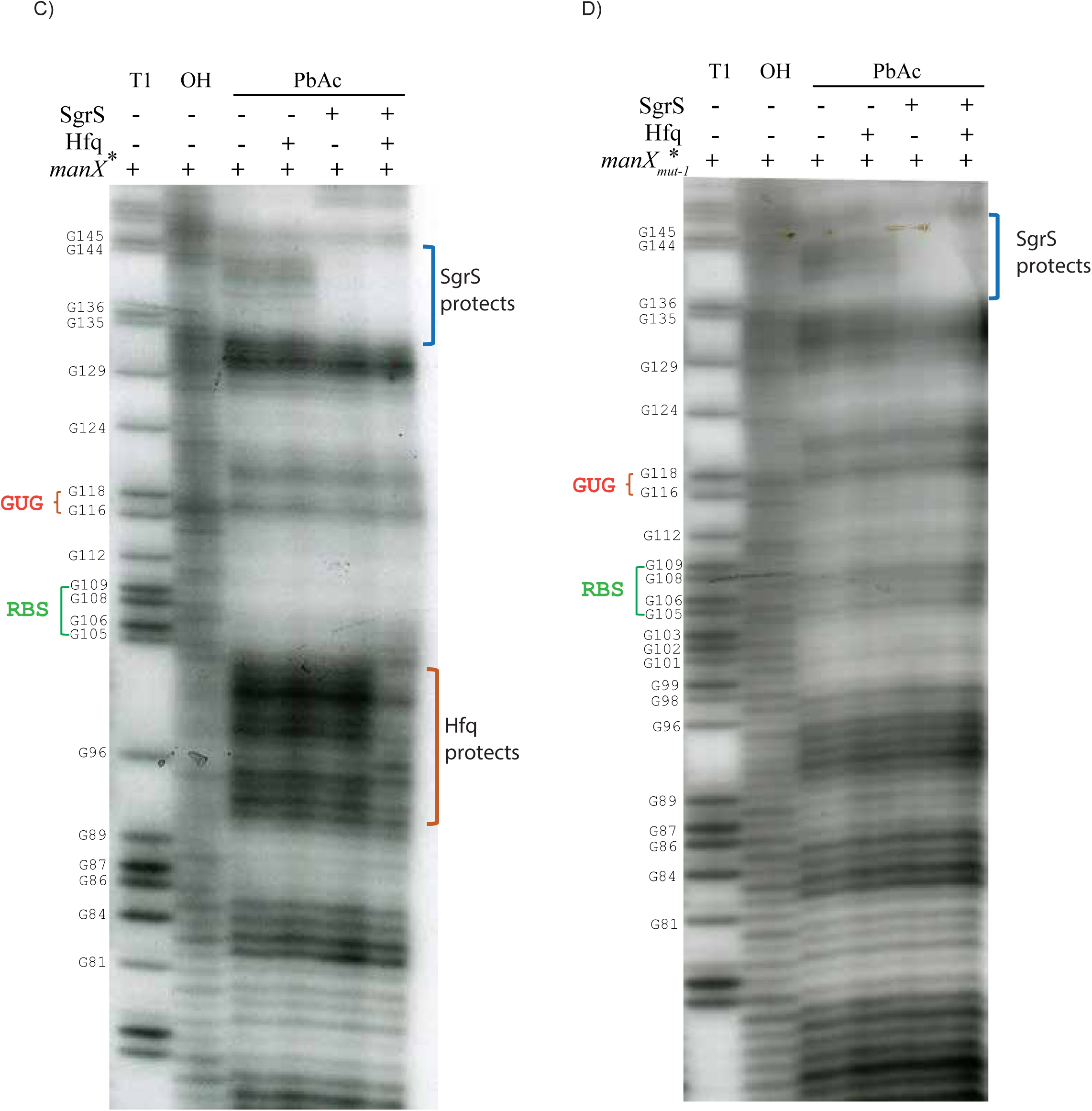
Footprinting maps SgrS, DicF and Hfq binding sites on *manX* RNA. In vitro transcribed *manX* mRNA containing the full-length 115-nt UTR and a portion of the coding region extending 51 nt downstream of the predicted DicF base pairing region was end labeled with ^32^P and incubated with and without unlabeled SgrS, DicF and Hfq to perform footprinting reactions. Samples were treated as follows: “T1,” RNase T1; “OH,” alkaline ladder; “PbAc,” lead acetate. Positions of G residues are indicated to the left of each gel image and nucleotides numbered as indicated in B. Positions of the GUG start codon and RBS are indicated to the left of each image. **A)** Footprinting SgrS and Hfq (left image) or DicF and Hfq (right image) binding sites on *manX* mRNA. **B)** Sequence and putative structure of *manX* following interaction with Hfq and SgrS or DicF. Positions of SgrS and DicF binding are indicated (from residues 139-167). The start codon is indicated by orange nucleotides. The Hfq binding site is highlighted in orange and the RBS is highlighted in green. **C-D)** Footprinting using reduced concentrations of Hfq (as described in Materials and Methods) in the absence and presence of SgrS. Wild-type *manX* with Hfq binding motif, AUAAUAAA is shown in C and the mut-1 Hfq binding motive, CGGCGGGA, is shown in D.

Experiments shown in Fig. 6A were performed with relatively high *manX* mRNA:Hfq ratios (1:37) and demonstrated Hfq binding to *manX* mRNA in the presence of sRNA. To test whether the sRNA promotes Hfq binding to *manX* mRNA when Hfq concentrations are limiting, we performed the footprint at lower *manX* mRNA:Hfq ratios (1:3) in the presence and absence of SgrS (Fig. 6C). At these lower ratios, we saw no protection of *manX* mRNA by Hfq in the absence of SgrS. In the presence of SgrS, we again observed protection by Hfq at the binding site adjacent to the RBS (Fig. 6C). As a control, we used the *manX mut-1* mRNA (Fig. 2B) and saw protection by SgrS, but no protection by Hfq, even in the presence of SgrS (Fig. 6D).

### *DicF is a weaker regulator of* manX

Compared to DicF, SgrS is a stronger repressor of *manX* translation (compare repression in Fig. 2B to Fig. 5A). Our data suggest that each of these sRNAs mediates translational regulation indirectly, via promoting Hfq binding to a site in the 5′ UTR adjacent to the RBS (Fig. 6A, C). To explore the basis for the different efficiencies of regulation, we conducted experiments to measure the affinity of sRNA-mRNA interactions and sRNA-Hfq interactions. We reasoned that differences in the binding affinity of the sRNAs *for manX* mRNA and/or Hfq, could be important determinants of regulatory efficiency for each sRNA-mRNA-Hfq interaction. Electrophoretic mobility shift assays (EMSAs) were used to measure the specific binding of SgrS and DicF individually to *manX* mRNA in vitro. We found that SgrS base paired with *manX* mRNA with a *K_D_* of 4.53 µM (Fig. 7A). In a previous study, we found that in vivo, the *K_D_* for SgrS binding to full-length *manXYZ* mRNA (with both *manX* and *manY* binding sites) was 2.3 µM (46). Thus, our in vitro measurement is in good agreement with the in vivo data for SgrS. DicF interacted less strongly with *manX* mRNA, with a *K_D_* of 21.8 µM (Fig. 7A).

**Figure 7.**
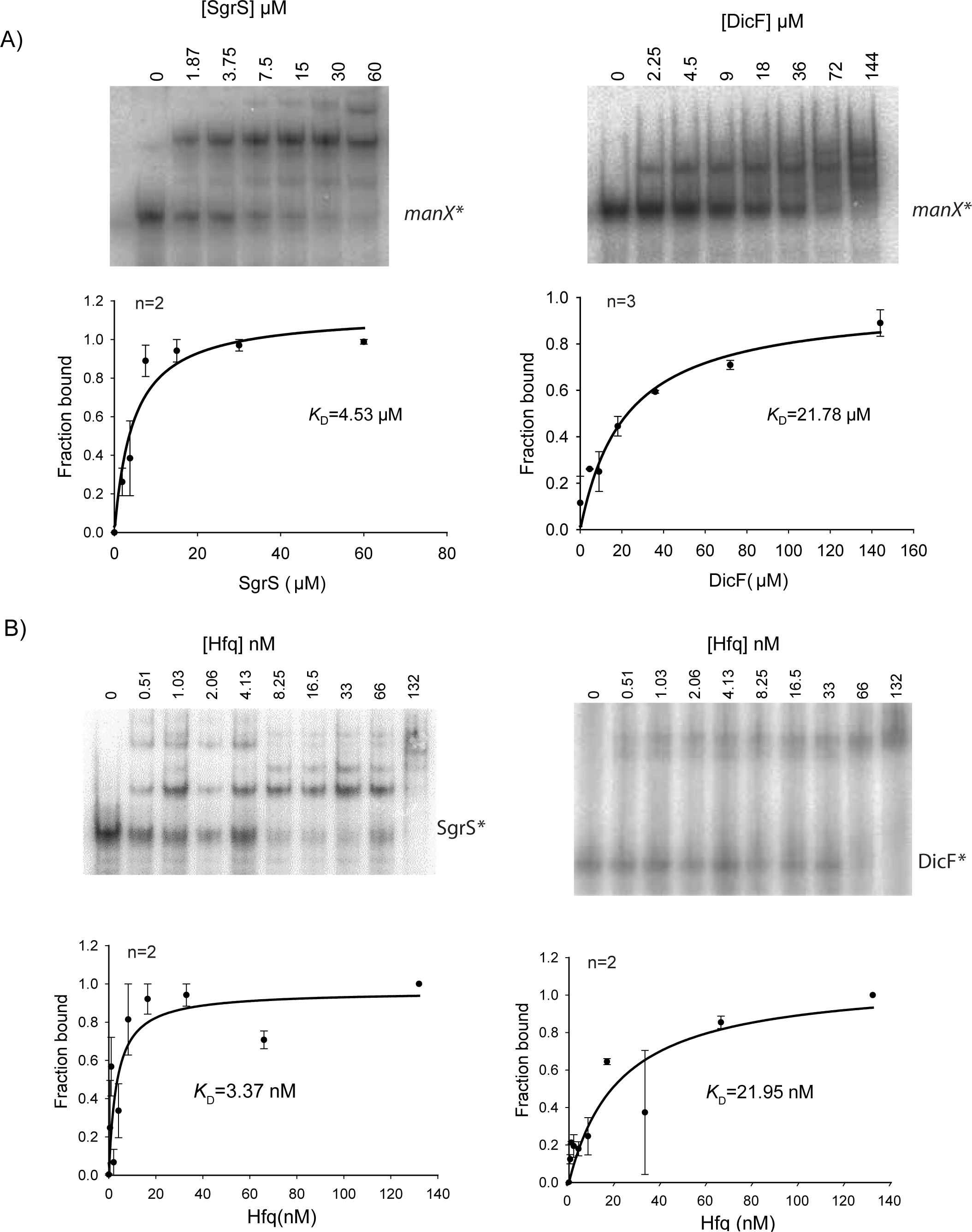
In vitro analyses of sRNA binding to *manX* mRNA and Hfq. **A)** Native gel electrophoresis was used to examine binding of *manX* mRNA with SgrS and DicF sRNAs. In vitro transcribed ^32^P-labeled *manX* mRNA (0.01 pmol) was mixed with indicated amounts of cold SgrS and incubated at 37^º^C for 30 min. The reaction mixture was resolved on a chilled native acrylamide gel. Bands were quantified and the fraction of *manX* mRNA bound was calculated and plotted to calculate *K*_D_. **B)** Gel mobility shift assay for SgrS (right) or DicF (left) and Hfq. Measured band densities (n replicates, top left) were plotted to determine the dissociation constants.

EMSAs to monitor interactions of each sRNA with Hfq also revealed differences between SgrS and DicF. The Hfq-SgrS interaction was relatively strong, with a calculated *K_D_* of 3.37 nM (Fig 7B). Hfq bound DicF less tightly with a calculated *K_D_* of 22.0 nM (Fig. 7B). The dissociation constant values we calculated for Hfq and SgrS or DicF are in a similar range as those reported previously for the binding of Hfq to OxyS, RyhB, DsrA, and Spot 42 sRNAs (11,47-49). Taken together, our data suggest that SgrS is a more efficient regulator of *manX* translation than DicF and that differences in sRNA-mRNA binding interactions and sRNA-Hfq interactions could influence the efficiency of regulation.

## DISCUSSION

Our study shows that two sRNAs, SgrS and DicF, base pair with *manX* mRNA at distinct sites in the coding region, outside the window that would allow translational repression via a direct ribosome occlusion mechanism. Instead, we propose that Hfq acts as the direct regulator of *manX* translation and that the sRNAs play an accessory or secondary role. This model is supported by multiple lines of evidence presented in this study. In vivo, SgrS cannot repress *manX* translation in the absence of Hfq (Fig. 1C). This is in contrast with regulation of another SgrS target, *ptsG*, which is known to occur via the canonical (direct) mechanism of translational repression (17,50). SgrS efficiently represses *ptsG* translation in an *hfq* mutant background (Fig. 1B). Loss of regulation by both sRNAs is seen in an *hfq^+^* background when the Hfq binding site upstream of the *manX* RBS is mutated (Figs. 2B, 5A). Structural analyses clearly demonstrate that SgrS and DicF bind to sites in the coding region of *manX* mRNA and have no impact on the structure near the TIR. In contrast, in the presence of either SgrS or DicF, Hfq binds to the same site on *manX* mRNA, directly adjacent to the RBS (Fig. 6A). (The Hfq binding site identified on *manX* mRNA in this study is consistent with Hfq-binding peaks in the same region identified in genome-wide Hfq crosslinking studies (51,52).) At low mRNA:Hfq ratios, the sRNA is required for Hfq binding to the RBS-adjacent site on *manX* mRNA (Fig. 6C). Differences in the relative strength of *manX* translational regulation promoted by SgrS and DicF were correlated with the strength of sRNA-mRNA and sRNA-Hfq interactions (Fig. 7A, B). Collectively, our data are consistent with a non-canonical mechanism of regulation where the sRNAs play a guide-like role in regulation by promoting Hfq binding to a site near the *manX* RBS so that Hfq itself directly interferes with ribosome binding (Fig. 8). This model contrasts with the canonical model of bacterial sRNA-mediated translational repression where sRNAs are the direct competitors of ribosome binding, while the chaperone Hfq assumes the secondary role. Here, the chaperone Hfq swaps its role with the RNA partner.

**Figure 8.**
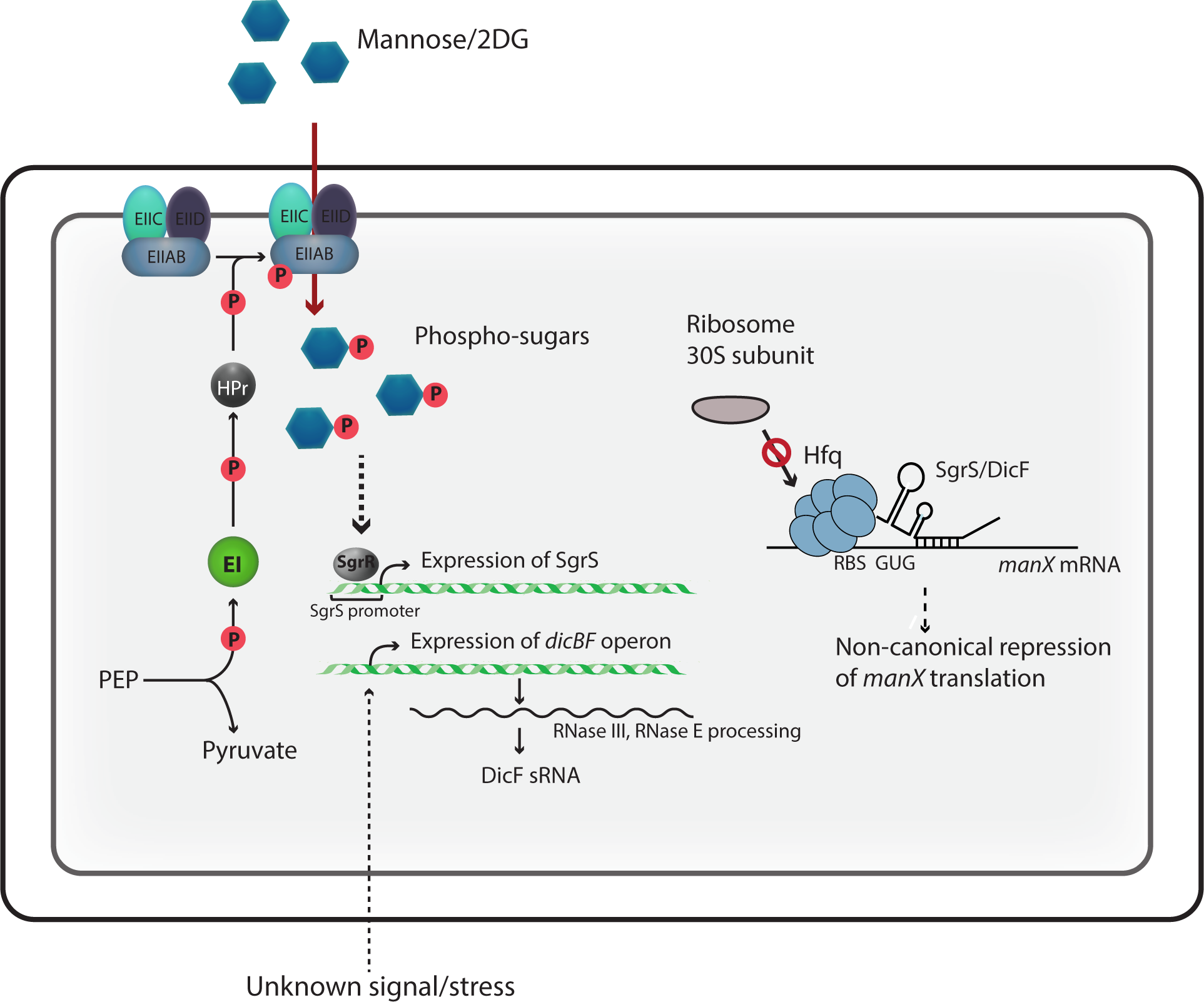
A model for the non-canonical roles played by two distinct sRNAs, SgrS and DicF, to repress *manX* translation via an Hfq-dependent mechanism.

The role of Hfq in post-transcriptional regulation of gene expression has been heavily studied, but much remains unclear. Bläsi and coworkers have studied several examples of Hfq-mediated regulation, including sRNA-dependent and ‐independent cases. Early work suggested that Hfq could directly regulate *ompA* mRNA (53), but this regulation was later shown to be mediated through the canonical mechanism of translational repression and dependent on the sRNA MicA (54). Hfq was also implicated in translational repression of *fur* mRNA (55) and translational autorepression of *hfq* mRNA (56), and so far there is no evidence that these effects require an sRNA. The Massé group discovered Hfq-dependent translational repression of two mRNAs, *shiA* (57) and *cirA* (58), and determined that the sRNA RyhB antagonizes Hfq-dependent repression to activate translation of both mRNAs. Chen and Gottesman recently discovered apparently direct translational repression by Hfq of *mutS* mRNA (59). The Hfq binding site on *mutS* mRNA is too far upstream of the TIR for direct occlusion of ribosome binding. Instead, Hfq binding appears to induce structural rearrangement of *mutS* mRNA in a way that inhibits translation.

To our knowledge, the only other example of sRNA-dependent regulation mediated by Hfq as the putative direct regulator is Spot 42 sRNA regulation of *sdhC* mRNA (28). Spot 42 was observed to bind far <u>upstream</u> of the *sdhC* RBS to carry out Hfq-dependent translational repression. For this sRNA-mRNA pair, the role of Spot 42 in regulation remains unclear. Spot 42 and Hfq are clearly required for regulation of *sdhC* in vivo, but in vitro, Hfq can bind efficiently to *sdhC* mRNA when present in a 1:1 mRNA:Hfq ratio, even in the absence of sRNA (Desnoyers and Masse, 2012). These findings contrast with our results, in several ways. For *manX* mRNA, the sRNA binding sites are located downstream of the TIR rather than upstream (Figs. 2B, 5B). Moreover, Hfq did not bind *manX* mRNA in vitro, even at a 1:3 mRNA:Hfq ratio unless SgrS was also present (Fig. 6C). Our results suggest that the sRNAs carry out the task of substrate recognition that subsequently allows the protein partner to be recruited to a binding site that is otherwise not efficiently or stably bound. Similar mechanisms are widely utilized by CRISPR guide RNAs, and eukaryotic non-coding RNAs, including small interfering RNAs, microRNAs, and small nucleolar RNAs. All of these types of RNAs act as a part of a ribonucleoprotein complex where the RNA component recognizes the substrate nucleic acid and promotes activity of the protein component at the correct site.

In bacteria, we have yet to uncover the mechanistic details of regulation carried out by the vast majority of sRNAs, but of those for which mechanisms have been established, sRNAs are typically the primary effectors of regulation. This raises some intriguing questions. Is the “canonical” mechanism of sRNA-mediated regulation with sRNA as primary regulator really the most common, or have computational and experimental approaches used to study sRNAs been biased toward discovery of these mechanisms because they were the first type described? Regardless of the prevalence of each of these two different mechanisms, what features distinguish them and make one or the other more favorable for regulation of a given mRNA target?

One advantage of sRNA-mediated regulation that involves base pairing interactions outside the TIR could be that it provides a larger and more diverse sequence space to evolve new regulatory interactions. We have found that regulation of *ptsG*, the primary glucose transporter, by SgrS is conserved among *E. coli* relatives where SgrS orthologs were found (60,61). The SgrS-*ptsG* mRNA interaction involves pairing between the most highly conserved seed region of SgrS and the *ptsG* RBS, a region where the sequence is highly conserved for ribosome binding. In contrast, SgrS-dependent regulation of *manX* involves a less well-conserved portion of SgrS and the coding sequence of *manX,* and this interaction is not entirely conserved among enteric species (21). Analyses by Peer and Margalit indicate that the SgrS-*ptsG* mRNA interaction evolved first, with the binding sites on both mRNA and sRNA co-appearing in evolutionary time (62). Their data suggest that SgrS-*manX* mRNA interaction evolved much later. So, SgrS first established a regulatory interaction with *ptsG* in an ancestral organism of the order *Enterobacteriales,* which established this sRNA regulator in the genome and allowed evolution of interactions with additional targets. Other recent work on sRNA evolution suggests similar target acquisition mechanisms where sRNAs establish one target and gradually establish other interactions with the concurrent evolution of Hfq (62,63). Perhaps flexibility in regulatory mechanisms, *e.g.,* where the sRNA can act as either a primary or accessory regulator along with Hfq, facilitates rapid evolution of additional sRNA-target interactions.

Regulation of *manX* translation by DicF, an sRNA encoded on the cryptic prophage Qin on the *E. coli* chromosome, was confirmed in this study. Like other small RNAs encoded on horizontally acquired genetic elements like prophages and pathogenicity islands, DicF is poorly characterized. However, research over the last decade, suggests that horizontally-acquired sRNAs are crucial regulators of bacterial physiology, growth, and stress responses (64-66). For instance, the sRNA InvR, encoded in *Salmonella* pathogenicity island 1, is a major regulator of outer membrane porin OmpD (67). IpeX, an sRNA encoded on the cryptic prophage DLP12 in *E. coli*, is a regulator of outer membrane porins, OmpC and OmpF (68). DicF was identified in the 1980s when it was observed to cause a filamentation phenotype when expressed from a multi-copy plasmid (69). We recently demonstrated that DicF directly regulates translation of mRNA targets encoding diverse products involved in cell division and metabolism (31). These include mRNAs encoding the tubulin homolog FtsZ, xylose uptake regulator XylR, and pyruvate kinase PykA (31,70). Our current study extends the DicF targetome to include *manX*. Though SgrS and DicF share a common target in *manX*, these sRNAs are not expressed under the same conditions. We did not see DicF expression when cells were challenged with αMG or 2DG (data not shown). Under standard laboratory growth conditions, the *dicBF* operon is not expressed, and we do not yet know the signal that triggers the expression of this operon. Further research aimed at uncovering the physiological conditions stimulating DicF production may provide insight into the biological role of DicF-mediated *manX* regulation.

A long-held notion about sRNA-mediated gene regulation in eukaryotes is that the primary role of sRNAs is target recognition, while the associated protein partners perform the primary regulatory function of gene silencing or translational repression. In bacteria, the prevailing model has been the oppositethat the sRNA is the primary regulator and associated proteins play secondary roles in promoting RNA stability or making the regulation irreversible (in the case of mechanisms involving mRNA degradation). Our findings, along with one other recent report on a similar non-canonical mechanism of regulation in bacteria (28) suggest that bacteria can utilize a broader range of sRNA-mediated regulatory strategies than previously suspected. So, while there are considerable differences among the domains in terms of the mechanisms of translation initiation, sites of sRNA binding, and the nature of the ribonucleoprotein complexes carrying out regulation, sRNA-directed recruitment of regulatory proteins to mRNA targets appears to be a common mode of regulation in all three domains of life.

## ACKNOWLEDGEMENTS

We thank members of the Vanderpool laboratory and the laboratory of James Slauch for productive discussions and Peter Orlean for sharing equipment important for carrying out experiments. Eric Massé and his lab members for advice and critical reading of the manuscript. We are grateful to Raya Romm, Hannah Margalit, Jai Tree, and David Tollervey for sharing their analyses and interpretation of Hfq and RNase E cross-linking datasets.

